# Locus Coeruleus Contrast and Diffusivity: Effects of Age and Relations to Memory

**DOI:** 10.1101/2024.03.25.586618

**Authors:** Ilana J. Bennett, Jason Langley, Andrew Sun, Kitzia Solis, Aaron R. Seitz, Xiaoping P. Hu

## Abstract

Neurocognitive aging researchers are increasingly focused on the locus coeruleus, a neuromodulatory brainstem structure that degrades with age. With this rapid growth, the field will benefit from consensus regarding which magnetic resonance imaging (MRI) metrics of locus coeruleus structure are most sensitive to age and cognition. To address this need, the current study acquired magnetization transfer- and diffusion-weighted MRI images in younger and older adults who also completed a free recall memory task. Results revealed significantly larger differences between younger and older adults for maximum than average magnetization transfer-weighted contrast (MTC), axial than mean or radial single-tensor diffusivity (DTI), and free than restricted multi-compartment diffusion (NODDI) metrics in the locus coeruleus; with maximum MTC being the best predictor of age group. Age effects for the MTC and NODDI metrics interacted with sex such that larger age group differences were seen in males than females. Age group differences were also larger for DTI metrics in the rostral, and NODDI metrics in the caudal, locus coeruleus subdivision. Within older adults, however, there were no significant effects of age on any measure of locus coeruleus structure. Finally, independent of age and sex, higher restricted diffusion in the locus coeruleus was significantly related to better (lower) recall variability, but not mean recall. Whereas MTC has been widely used in the literature, our comparison between the average and maximum MTC metrics, and inclusion of DTI and NODDI metrics, make important and novel contributions to our understanding of the aging of locus coeruleus structure.

## Introduction

The locus coeruleus has garnered significant attention in neurocognitive aging research in recent years given that it is one of the first brain regions to accumulate tau pathology implicated in Alzheimer’s Disease (Braak et al., 2011; Jacobs et al., 2021). Even in the absence of disease, histopathological studies in humans have found reductions in locus coeruleus neurons with age (Flood & Coleman, 1988; German et al., 1988; Manaye et al., 1995; Vijayashankar & Brody, 1979) c.f. (Mouton et al., 1994). These noradrenergic neurons project throughout the brain (Foote et al., 1983; Gatter & Powell, 1977; Waterhouse et al., 1983) and are thought to play a neuromodulatory role in a broad range of cognitive processes (Berridge & Waterhouse, 2003; Dahl, Kulesza, et al., 2023; Mather & Harley, 2016; Poe et al., 2020; Sara, 2009), including memory. Whereas recent advances in magnetic resonance imaging (MRI) have made it possible to reliably image and segment this small brainstem structure *in vivo* (Priovoulos et al., 2018; Yi et al., 2023), this rapidly growing field will benefit from consensus regarding which metrics of locus coeruleus structure are most sensitive to age and cognitive performance.

Structural “integrity” of the locus coeruleus is most often assessed using either fast spin-echo T1-weighted or magnetization transfer-weighted MRI sequences in which signal intensity (or contrast) is thought to be driven by the magnetic properties of neuromelanin (Betts et al., 2019; Nakane et al., 2008; Priovoulos et al., 2018; Trujillo et al., 2019, 2023) c.f. (Watanabe, 2023), which is a pigmented byproduct of norepinephrine synthesis in the locus coeruleus. Magnetization transfer contrast (MTC) ratio in the locus coeruleus is measured relative to the pons and then either averaged across voxels within an anatomical mask of the locus coeruleus (e.g., (Liu et al., 2019) or extracted from the voxel(s) with the maximum value (e.g., (Clewett et al., 2016; Jacobs et al., 2021; Takahashi et al., 2015). However, concerns have been raised about whether maximum MTC accurately captures the locus coeruleus (Liu et al., 2017) as age-related neuronal loss may be unequally distributed across the structure, comparable to what is seen in Alzheimer’s Disease (German et al., 1992).

Using these MRI approaches, cross-sectional studies have consistently found a quadratic or inverted U-shaped function in locus coeruleus MTC across the adult lifespan. Locus coeruleus MTC increases with age from 20 to ∼60 years and decreases with age after ∼60 years old (Eckert et al., 2023; Jacobs et al., 2021; Liu et al., 2019; Shibata et al., 2006), although some studies within only older adults have found no effect of age on locus coeruleus MTC (Calarco et al., 2022; Giorgi et al., 2022). Results have been more mixed when comparing extreme age groups, with some studies reporting higher locus coeruleus MTC in older than younger adults (Clewett et al., 2016; Pievani et al., 2020), and others finding no age group differences (Dahl, Bachman, et al., 2023; Dahl et al., 2019; Hämmerer et al., 2018; Porat et al., 2022). There is some evidence that these age effects are more prominent in the rostral than caudal subdivision of the locus coeruleus (Betts et al., 2017; Dahl et al., 2019; Liu et al., 2019, 2020), consistent with the rostral subdivision being more vulnerable to cell loss in aging, similar to what is seen in Alzheimer’s Disease (German et al., 1992). Of the handful of studies that have examined sex difference in locus coeruleus MTC, most have reported no significant sex effects (Calarco et al., 2022; Jacobs et al., 2021; Liu et al., 2019; Shibata et al., 2006), whereas at least one observed lower locus coeruleus MTC in females than males that was independent of age group (Clewett et al., 2016).

In contrast to MTC, there has been a dearth of literature using other approaches that can assess microstructural “integrity” of the locus coeruleus, such as diffusion-weighted MRI. When diffusion of molecular water in each voxel is modelled using a single-tensor, diffusion tensor imaging (DTI) metrics such as mean diffusivity (MD) measure the average rate of diffusion, which varies with tissue properties including neurodegeneration, neuroinflammation, and an accumulation of pathology. A small group of recent studies have revealed lower diffusivity in older than younger adults in locus coeruleus gray matter (Langley et al., 2020, 2022; Porat et al., 2022) c.f. (Pievani et al., 2020). To date, however, no studies have assessed effects of age on locus coeruleus using multi-compartment diffusion approaches, such as Neurite Orientation Dispersion and Density Imaging (NODDI; (Zhang et al., 2012). NODDI may be better suited than DTI for capturing microstructural properties in gray matter as it models compartments of diffusion that are invariant to tissue organization, yielding stronger effects of age and cognition in regions such as the hippocampus (Venkatesh et al., 2020). Moreover, although some of these studies assessed locus coeruleus using both DTI and MTC (Pievani et al., 2020; Porat et al., 2022), they did not compare the sensitivity of these imaging modalities to age.

Age-related degradation of the locus coeruleus, as measured by either MTC or diffusion DTI, NODDI), would have significant consequences for cognitive processes mediated by brain regions innervated by its noradrenergic projections, such as memory. Consistent with this view, previous studies have shown that lower MTC (Dahl et al., 2019; Dahl, Kulesza, et al., 2023; Elman et al., 2021; Hämmerer et al., 2018; Jacobs et al., 2021; Solders et al., 2020) and lower DTI diffusivity (Langley et al., 2020, 2022) in locus coeruleus gray matter relates to worse memory performance in older adults. For diffusivity, these relationships have been observed in both rostral and caudal locus coeruleus (Langley et al., 2022). Whereas these studies have focused on average memory performance (e.g., mean across trials), a measure of variability between trials may be more sensitive to locus coeruleus structure as any degradation may result in momentary disruptions to attention via locus coeruleus neuromodulation, resulting in less consistent performance (Unsworth & Robison, 2017).

The current study sought to address the gaps of prior work by acquiring magnetization transfer- and diffusion-weighted MRI images in younger and older adults who also completed a word list free recall memory task. Effects of age on locus coeruleus structure were assessed by comparing extreme age groups (younger, older) as well as relationships to age within older adults for MTC (average, maximum), DTI (mean diffusivity, MD; axial diffusivity, AD; radial diffusivity, RD), and NODDI (restricted, free) metrics. Effects of sex (male, female) and subdivision (rostral, caudal) on aging of locus coeruleus structure were also examined, as were relationships between locus coeruleus structure and memory performance using measures of both mean recall and recall variability.

## Methods

### Participants

Fifty-two younger and 77 older adults who were recruited from the University of California, Riverside and surrounding communities met our inclusion criteria, which included having normal general cognition (e.g., <17 on the telephone Montreal Cognitive Assessment, MoCA)(Pendlebury et al., 2013), self-reported absence of major health conditions (e.g., stroke, dementia, diabetes); and being free of conditions that would prevent them from being able to enter the MRI scanner (e.g., non-MR compliant implants, difficulty lying in the supine position, claustrophobia). One younger adult was excluded after data collection due to uncorrectable mis-registration that yielded inaccurate MRI metrics. Demographic and neuropsychological data for the final sample of 51 younger (18-26 years) and 77 older (60-87 years) adults are provided in Table 1.

**Table 1.** Demographic and neuropsychological test data.

| | Younger Adults | Older Adults | $t / \chi^2$ |
| --- | --- | --- | --- |
| N | 51 | 77 | n/a |
| Age (years) | 20.2 ± 2.2 | 69.2 ± 5.9 | **54.2** |
| Sex (% female) | 53.1% | 66.2% | 2.1 |
| Education (years) | 13.7 ± 1.5 | 15.6 ± 2.7 | **4.3** |
| Ethnicity (% Non-Hispanic) | 64.7% | 71.7% | 3.3 |
| Race (% White) | 23.9% | 73.6% | **28.1** |
| Handedness (% right-handed) | 93.5% | 100.0% | **7.2** |
| MoCA | n/a | 19.9 ± 1.5 | n/a |
| Free recall mean | 5.1 ± 1.2 | 4.8 ± 1.4 | -0.9 |
| Free recall variability | 0.2 ± 0.1 | 0.3 ± 0.2 | 1.2 |

*Notes.* Mean (mean and standard deviation, $M \pm SD$ ) or percent (%) scores are provided for each age group. Significant group differences at $p < 0.05$ are indicated by bolded $t$ (mean scores) or $\chi^2$ (% scores) statistics. Excludes all or partial demographic (5 younger, 3 older), telephone Montreal Cognitive Assessment (MoCA; maximum score 22; 3 older), and free recall (3 younger, 9 older) data because responses were not recorded or tasks were not completed.

All participants provided informed consent and received course credit or financial compensation for participation. Study procedures were approved by the Institutional Review Board of the University of California, Riverside.

### Magnetic Resonance Imaging Data

#### Acquisition

Imaging data were acquired on a 3T MRI scanner (Prisma, Siemens Healthineers, Malvern, PA) at the Center for Advanced Neuroimaging at the University of California, Riverside. Excitation was performed using the body coil on the scanner and signal was received using a 32-channel receive only coil.

A T_1_-weighted MP-RAGE image was acquired with the following parameters: echo time (TE)/repetition time (TR)/inversion time = 3.02/2600/800 ms, GRAPPA acceleration factor = 2, flip angle = 8°, voxel size = 0.8×0.8×0.8 mm^3^.

Magnetization transfer-prepared gradient echo (MT-GRE) images were acquired with the following parameters: TE/TR = 3.21/385 ms, flip angle = 40°, field of view (FOV) = 220×186 mm^2^, matrix size = 512×432, slice thickness = 3 mm, magnetization transfer preparation pulse (flip angle = 370°, 1.5 kHz off-resonance, duration =10 ms), 4 averages. Slices were positioned perpendicular to the dorsal edge of the brainstem at the midline along the fourth ventricle.

Diffusion-weighted single-shot spin-echo, echo planar images were acquired with the following parameters: TE/TR = 75/4100 ms, FOV = 202×170 mm^2^, matrix size = 176×148, voxel size = 1.15×1.15×2.5 mm^3^, and 32 slices with no gap. Slices were aligned parallel to the hippocampus and covered the brain from the middle of the cerebellum to the striatum. Monopolar diffusion-encoding gradients were applied in 30 directions with *b* values of 500 s/mm^2^ and 2000 s/mm^2^. Two sets of *b*=0 images were acquired, with the two sets having opposite polarities of phase-encoding direction for the correction of susceptibility distortion (Andersson et al., 2003).

#### Regions of interest

A standardized atlas was used to define bilateral locus coeruleus, as well as its rostral and caudal subdivisions (Langley et al., 2020). A rectangular midline reference region was defined in the pons (Figure 1). For each participant, regions of interest were aligned from Montreal Neurological Institute (MNI) space to their MT-GRE and diffusion-weighted space using a transformation that concatenated an alignment between MNI T_1_-weighted space and their T_1_-weighted image using FMRIB’s Linear Image Registration Tool (FLIRT) and FMRIB’s Nonlinear Image Registration Tool (FNIRT) in the FMRIB Software Library (FSL)(Jenkinson et al., 2002) and an alignment between the participant’s T_1_-weighted and either their MT-GRE (using the averaged MT-GRE image) or diffusion-weighted (using the average *b*=0 image) space using separate rigid body transformations with a boundary-based registration cost function. Each aligned region of interest was thresholded at 50% and binarized.

**Figure 1.**
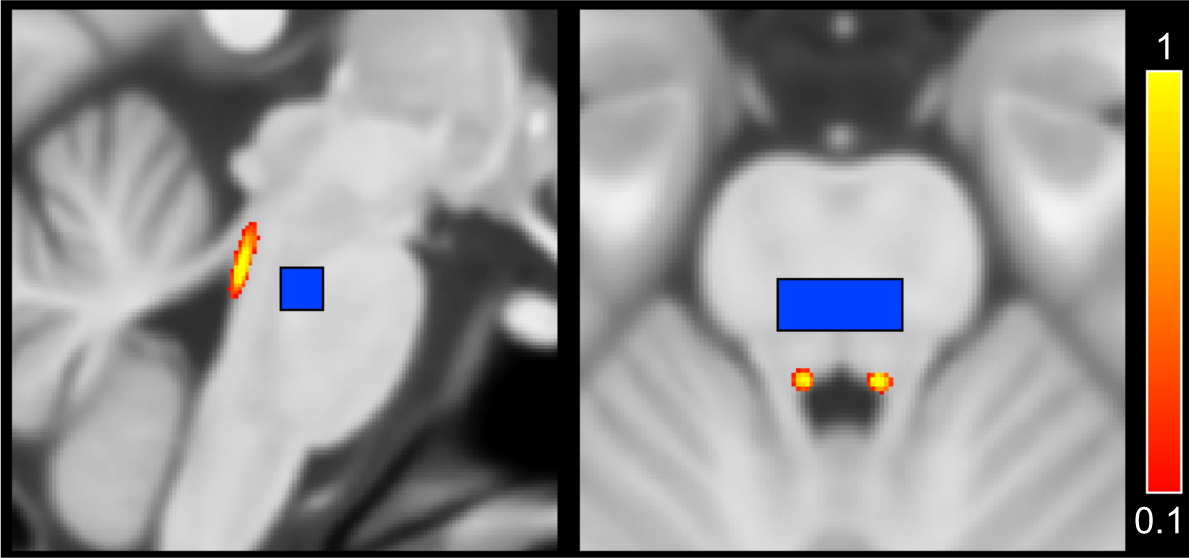
Sagittal (left) and axial (right) views of the atlas-based bilateral locus coeruleus (red-yellow) and rectangular midline pontine reference region (blue) in Montreal Neurological Institute (MNI) T_1_-weighted space.

#### MTC metrics

For each participant, individual measurements from the MT-GRE acquisition were corrected for motion by registering them to the first image using a FLIRT rigid-body transformation and then averaged. Finally, a magnetization transfer contrast (MTC) image was calculated using the following equation:

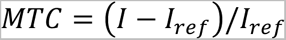

where *I* denotes the intensity of a voxel in the MT-GRE image and *I_ref_* is the mean intensity of the pontine reference region. Average and maximum MTC metrics were calculated by averaging MTC values or finding the peak MTC value within the MT-GRE-aligned locus coeruleus regions of interest, respectively. Individual values were excluded from the MTC analyses if they exceeded four standard deviations from the sample mean (2 older).

#### Diffusion metrics

For each participant, raw diffusion-weighted data were corrected for motion, susceptibility distortions, and eddy currents using FSL’s TOPUP and EDDY (Andersson et al., 2003; Andersson & Sotiropoulos, 2016). Diffusion tensor imaging (DTI) metrics (mean diffusivity, MD; axial diffusivity, AD; radial diffusivity, RD) were estimate using FSL’s DTIFIT. Neurite orientation dispersion and density imaging (NODDI) metrics (free diffusion, also known as ‘fiso’; restricted diffusion, also known as ‘ficvf’) were estimated using the NODDI toolbox v1.0.1 in MATLAB (Zhang et al., 2012). Mean diffusion metrics were calculated by averaging values within the diffusion-aligned locus coeruleus regions of interest. Eight participants (3 younger, 5 older) were excluded from all NODDI analyses for having free diffusion values of zero, indicating poor model fit in the first stage. Individual values were also excluded from the DTI (1 younger) and NODDI (2 younger) analyses if they exceeded four standard deviations from the sample mean.

### Memory Test

Participants completed three unique trials of a free recall task. On each trial, participants repeat out loud each of 10 words that were shown one at a time on a computer screen (30 seconds total), followed by an immediate free recall test (90 seconds). Words were selected from the Auditory Verbal Learning Test (Rey, 1964) and did not repeat across trials. Mean recall was calculated as the average number of words recalled across trials. Recall variability (coefficient of variation) was calculated as the standard deviation across trials divided by the mean (Murtha et al., 2002). Twelve participants (3 younger, 9 older) were excluded from these analyses because they did not complete the memory task.

## Results

### Locus Coeruleus Structure Differs Between Younger and Older Adults

#### MTC Metrics

An Age Group (younger, older) × Sex (male, female) × Metric (average, maximum) mixed factorial ANOVA was first conducted using MTC values from the whole locus coeruleus. Next, an Age Group × Sex × Subdivision (rostral, caudal) mixed factorial ANOVA was conducted using only the average metric from the locus coeruleus subdivisions, excluding the peak MTC as it appeared in just one subdivision for each participant. Results from both analyses (whole, subdivision) are described below and presented in Tables 2-4 and Figure 2.

**Figure 2.**
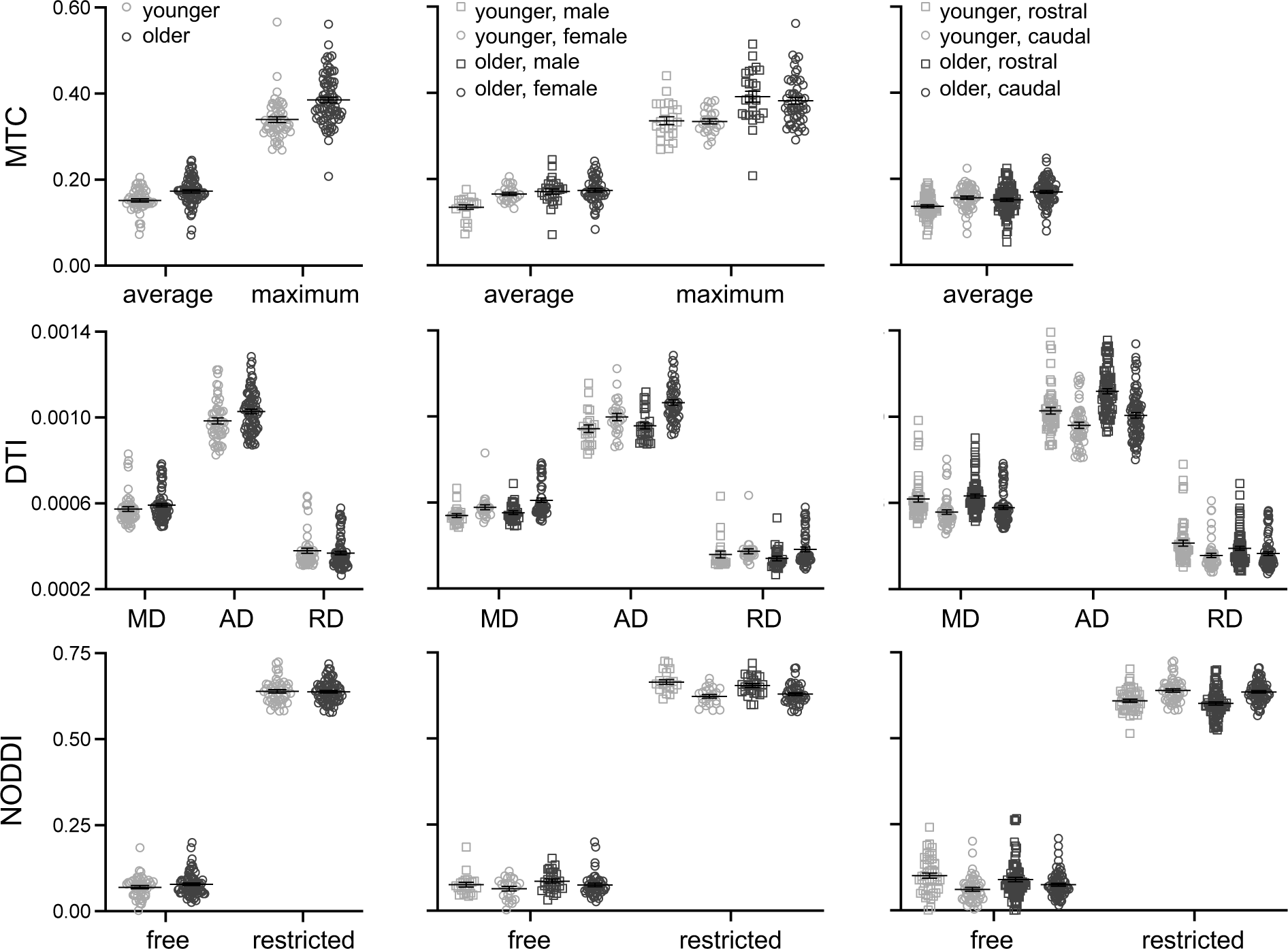
Age group differences in locus coeruleus structure are shown for each imaging metric (left column), with additional breakdowns by sex (middle column) and locus coeruleus subdivision (right column).

**Table 2.**
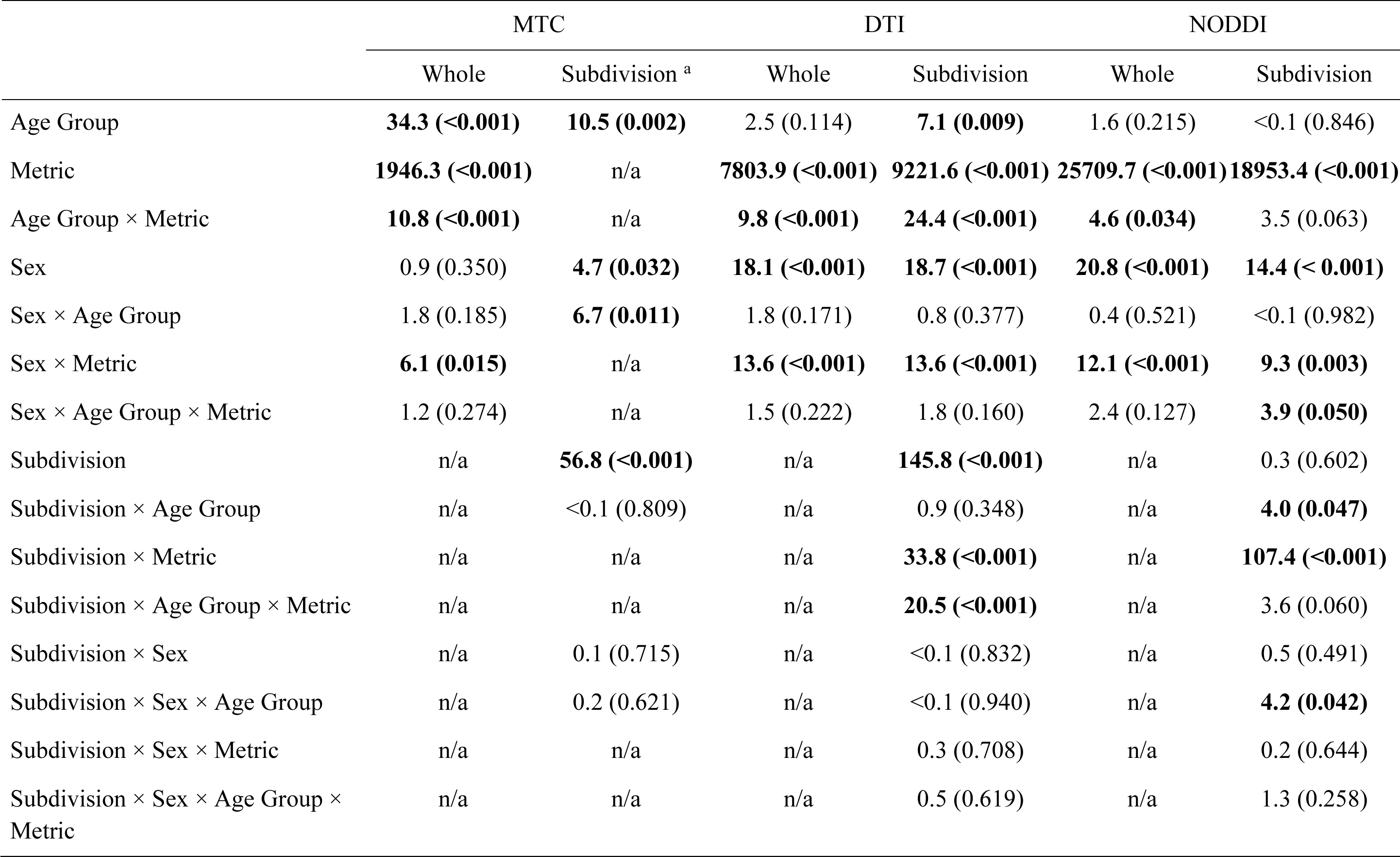
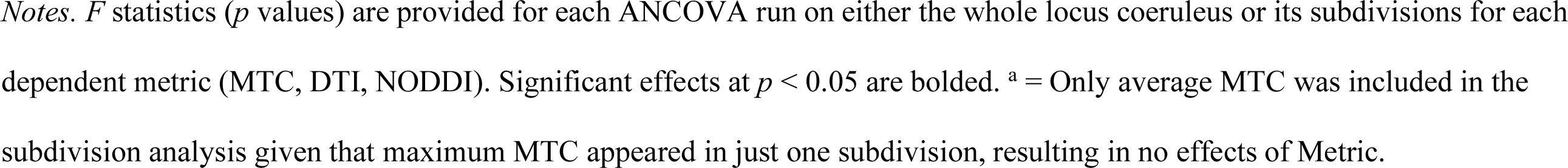
ANCOVA Results.

|  | MTC |  | DTI |  | NODDI |  |
| --- | --- | --- | --- | --- | --- | --- |
|  | Whole | Subdivision <sup>a</sup> | Whole | Subdivision | Whole | Subdivision |
| Age Group | **34.3 (<0.001)** | **10.5 (0.002)** | 2.5 (0.114) | **7.1 (0.009)** | 1.6 (0.215) | <0.1 (0.846) |
| Metric | **1946.3 (<0.001)** | n/a | **7803.9 (<0.001)** | **9221.6 (<0.001)** | **25709.7 (<0.001)** | **18953.4 (<0.001)** |
| Age Group × Metric | **10.8 (<0.001)** | n/a | **9.8 (<0.001)** | **24.4 (<0.001)** | **4.6 (0.034)** | 3.5 (0.063) |
| Sex | 0.9 (0.350) | **4.7 (0.032)** | **18.1 (<0.001)** | **18.7 (<0.001)** | **20.8 (<0.001)** | **14.4 (<0.001)** |
| Sex × Age Group | 1.8 (0.185) | **6.7 (0.011)** | 1.8 (0.171) | 0.8 (0.377) | 0.4 (0.521) | <0.1 (0.982) |
| Sex × Metric | **6.1 (0.015)** | n/a | **13.6 (<0.001)** | **13.6 (<0.001)** | **12.1 (<0.001)** | **9.3 (0.003)** |
| Sex × Age Group × Metric | 1.2 (0.274) | n/a | 1.5 (0.222) | 1.8 (0.160) | 2.4 (0.127) | **3.9 (0.050)** |
| Subdivision | n/a | **56.8 (<0.001)** | n/a | **145.8 (<0.001)** | n/a | 0.3 (0.602) |
| Subdivision × Age Group | n/a | <0.1 (0.809) | n/a | 0.9 (0.348) | n/a | **4.0 (0.047)** |
| Subdivision × Metric | n/a | n/a | n/a | **33.8 (<0.001)** | n/a | **107.4 (<0.001)** |
| Subdivision × Age Group × Metric | n/a | n/a | n/a | **20.5 (<0.001)** | n/a | 3.6 (0.060) |
| Subdivision × Sex | n/a | 0.1 (0.715) | n/a | <0.1 (0.832) | n/a | 0.5 (0.491) |
| Subdivision × Sex × Age Group | n/a | 0.2 (0.621) | n/a | <0.1 (0.940) | n/a | **4.2 (0.042)** |
| Subdivision × Sex × Metric | n/a | n/a | n/a | 0.3 (0.708) | n/a | 0.2 (0.644) |
| Subdivision × Sex × Age Group × Metric | n/a | n/a | n/a | 0.5 (0.619) | n/a | 1.3 (0.258) |

*Notes.* *F* statistics (*p* values) are provided for each ANCOVA run on either the whole locus coeruleus or its subdivisions for each dependent metric (MTC, DTI, NODDI). Significant effects at $p < 0.05$ are bolded. <sup>a</sup> = Only average MTC was included in the subdivision analysis given that maximum MTC appeared in just one subdivision, resulting in no effects of Metric.

**Table 3.** Whole Locus Coeruleus Descriptive Statistics.

|  | Younger |  | Older |  |
| --- | --- | --- | --- | --- |
|  | Male | Female | Male | Female |
| *MTC metrics* |  |  |  |  |
| Average | 0.14 ± 0.006 | 0.17 ± 0.006 | 0.17 ± 0.006 | 0.18 ± 0.004 |
| Maximum | 0.34 ± 0.011 | 0.33 ± 0.010 | 0.39 ± 0.010 | 0.38 ± 0.007 |
| *DTI metrics*<sup>u</sup> |  |  |  |  |
| MD | 0.54 ± 0.013 | 0.58 ± 0.012 | 0.55 ± 0.012 | 0.61 ± 0.009 |
| AD | 0.94 ± 0.018 | 1.00 ± 0.017 | 0.96 ± 0.016 | 1.06 ± 0.012 |
| RD | 0.36 ± 0.015 | 0.37 ± 0.014 | 0.34 ± 0.014 | 0.38 ± 0.010 |
| *NODDI metrics* |  |  |  |  |
| Restricted | 0.66 ± 0.006 | 0.62 ± 0.006 | 0.65 ± 0.006 | 0.63 ± 0.004 |
| Free | 0.07 ± 0.007 | 0.06 ± 0.006 | 0.09 ± 0.006 | 0.08 ± 0.005 |

*Notes.* Mean ± standard error are provided for each dependent metric (MTC, DTI, NODDI) in
the whole locus coeruleus, separately for each age and sex group. <sup>u</sup> = units × 10<sup>-3</sup>.

**Table 4.** Locus Coeruleus Subdivision Descriptive Statistics.

|  | Younger |  |  |  | Older |  |  |  |
| --- | --- | --- | --- | --- | --- | --- | --- | --- |
|  | Male |  | Female |  | Male |  | Female |  |
|  | Rostral | Caudal | Rostral | Caudal | Rostral | Caudal | Rostral | Caudal |
| *MTC metrics* |  |  |  |  |  |  |  |  |
| Average | 0.12 ± 0.006 | 0.15 ± 0.006 | 0.15 ± 0.006 | 0.17 ± 0.006 | 0.15 ± 0.006 | 0.17 ± 0.005 | 0.15 ± 0.004 | 0.17 ± 0.004 |
| *DTI metrics*<sup>u</sup> |  |  |  |  |  |  |  |  |
| MD | 0.58 ± 0.015 | 0.53 ± 0.014 | 0.62 ± 0.014 | 0.56 ± 0.013 | 0.60 ± 0.014 | 0.54 ± 0.013 | 0.65 ± 0.010 | 0.60 ± 0.009 |
| AD | 0.98 ± 0.019 | 0.91 ± 0.020 | 1.03 ± 0.018 | 0.98 ± 0.019 | 1.05 ± 0.018 | 0.94 ± 0.019 | 1.15 ± 0.013 | 1.04 ± 0.013 |
| RD | 0.38 ± 0.016 | 0.33 ± 0.014 | 0.41 ± 0.015 | 0.36 ± 0.013 | 0.37 ± 0.015 | 0.34 ± 0.013 | 0.40 ± 0.010 | 0.38 ± 0.009 |
| *NODDI metrics* |  |  |  |  |  |  |  |  |
| Restricted | 0.63 ± 0.008 | 0.66 ± 0.006 | 0.60 ± 0.007 | 0.62 ± 0.006 | 0.62 ± 0.007 | 0.65 ± 0.006 | 0.59 ± 0.005 | 0.63 ± 0.004 |
| Free | 0.09 ± 0.012 | 0.06 ± 0.007 | 0.10 ± 0.011 | 0.05 ± 0.007 | 0.10 ± 0.010 | 0.08 ± 0.007 | 0.08 ± 0.008 | 0.07 ± 0.005 |

*Notes.* Mean ± standard error are provided for each dependent metric (MTC, DTI, NODDI) in each locus coeruleus subdivision,
separately for each age and sex group. <sup>u</sup> = units × 10<sup>-3</sup>.

##### Age group differences

The whole locus coeruleus analysis revealed significant effects of Age Group and Metric, with higher MTC in older than younger adults and for the maximum than average metric. A significant Age Group × Metric interaction revealed that the age group difference was larger for the maximum (mean difference = M_diff_: 0.052) than average (M_diff_: 0.023) metric.

##### Age group differences varied by sex

The whole locus coeruleus analysis revealed a significant Sex × Metric interaction, with significantly higher MTC in females than males for the average (M_diff_: 0.017), but not maximum (M_diff_: -0.005), metric.

The locus coeruleus subdivision analysis further revealed significant effects of Sex and Sex × Age Group for the average MTC metric, with higher average MTC in females than males and higher average MTC in older than younger males (M_diff_: 0.028), but not females (M_diff_: 0.003).

##### Age group differences did not vary by subdivision

The locus coeruleus subdivision analysis revealed a significant main effect of Subdivision, with higher average MTC in the caudal than rostral subdivision. However, there were no significant interactions between Subdivision and Age Group.

#### DTI Metrics

An Age Group (younger, older) × Sex (male, female) × Metric (MD, AD, RD) mixed factorial ANOVA was conducted using DTI metrics from the whole locus coeruleus. Next, an Age Group × Sex × Metric × Subdivision (rostral, caudal) mixed factorial ANOVA was conducted using DTI metrics from the locus coeruleus subdivisions. Results from the latter are described below and results from both analyses (whole, subdivision) are presented in Tables 2-4 and Figure 2.

##### Age group differences

Results revealed significant effects of Age Group and Metric, with higher diffusivity in older than younger adults and for AD, then MD, and then RD. A significant Age Group × Metric interaction revealed that the age group difference was significant for AD (M_diff_: 0.069 ×10^-3^) and MD (M_diff_: 0.025 ×10^-3^), but not RD (M_diff_: 0.003 ×10^-3^).

##### Age group differences did not vary by sex

Results revealed significant effects of Sex and Sex × Metric with higher diffusivity in females than males and this sex difference was larger for AD (M_diff_: 0.080 ×10^-3^) than MD (M_diff_: 0.047 ×10^-3^) or RD (M_diff_: 0.031 ×10^-3^). However, there were no significant interactions between Sex and Age Group.

##### Age group differences varied by subdivision

Results revealed significant effects of Subdivision, Subdivision × Metric, and Subdivision × Age Group × Metric. The three-way interaction was probed using separate independent sample *t*-tests for each metric in each subdivision, which revealed that the age group difference was larger for rostral (M_diff_: 0.091 ×10^-3^; *p* < 0.001) than caudal (M_diff_: 0.046 ×10^-3^; *p* = 0.018) AD, but not significant for MD (rostral M_diff_: 0.014 ×10^-3^; caudal M_diff_: 0.021 ×10^-3^; *p*s > 0.11) or RD (rostral M_diff_: -0.024 ×10^-3^; caudal M_diff_: 0.009 ×10^-3^; *p*s > 0.13).

#### NODDI Metrics

An Age Group (younger, older) × Sex (male, female) × Metric (restricted, free) mixed factorial ANOVA was conducted using NODDI metrics from the whole locus coeruleus. Next, an Age Group × Sex × Metric × Subdivision (rostral, caudal) mixed factorial ANOVA was conducted using NODDI metrics from the locus coeruleus subdivisions. Results from both analyses (whole, subdivision) are described below and presented in Tables 2-4 and Figure 2.

##### Age group differences

The whole locus coeruleus analysis revealed a significant main effect of Metric, with higher restricted than free diffusion. A significant Age Group × Metric interaction revealed higher diffusion in older than younger adults for the free (M_diff_: 0.013), but not restricted (M_diff_: -0.002), metric. The Age Group × Metric interaction was not significant in the locus coeruleus subdivision analysis.

##### Age group differences varied by sex

The whole locus coeruleus analysis revealed significant Sex and Sex × Metric effects, with higher diffusion in males than females and this sex difference was significant for restricted (M_diff_: 0.033), but not free (M_diff_: 0.009), diffusion.

The locus coeruleus subdivision analysis further revealed a significant Sex × Age Group × Metric interaction, which was probed using separate independent sample *t*-tests for each metric in each sex group. Results revealed a non-significant trend for higher restricted diffusion in younger than older males (M_diff_: 0.014; *p* = 0.097) and higher free diffusion in older than younger males (M_diff_: 0.016; *p* = 0.110), but these effects in females did not approach significance (restricted M_diff_: <0.001; free M_diff_: 0.001; *p*s > 0.86).

##### Age group differences varied by subdivision

The locus coeruleus subdivision analysis revealed significant effects of Subdivision × Age Group, Subdivision × Metric, and Subdivision × Sex × Age Group. The three-way interaction was probed using separate independent sample *t*-tests for each sex group in each subdivision, which revealed significantly higher diffusion in older than younger females in the caudal (M_diff_: 0.014; *p* = 0.023), but not rostral (M_diff_: -0.012; *p* = 0.165), subdivision or in males in either subdivision (caudal M_diff_: 0.001; rostral M_diff_: 0.001; *p*s > 0.90) subdivision.

#### Best Predictor of Age Group

A stepwise logistic regression was conducted using all metrics from the whole locus coeruleus and sex as predictor variables entered with Forward Wald selection. Results revealed that maximum MTC was the best predictor of age group, *x*^2^(1) = 23.7, *p* < 0.001, correctly classifying 73.5% of participants. Classification accuracy was slightly improved when adding MD and RD (74.3%); AD and RD (75.2%); or MD, AD, and RD (77.0%) to the model.

When the stepwise logistic regression was conducted using all metrics from the locus coeruleus subdivisions and sex as predictor variables, results revealed that rostral AD was the best predictor of age group, *x*^2^(1) = 28.1, *p* < 0.001, correctly classifying 70.4% of participants. Classification accuracy was slightly improved when adding rostral RD (77.4%); rostral RD and caudal free (81.7%); or rostral RD, caudal free, and caudal average MTC (80.0%) to the model.

### Locus Coeruleus Structure Does Not Differ Within Older Adults

Within older adults, separate multiple regression analyses were conducted for each metric from each imaging modality (MTC: average, maximum; DTI: MD, AD, RD; NODDI: restricted, free), first in the whole locus coeruleus and then in each locus coeruleus subdivision.

Chronological Age, Sex, and their interaction were predictor variables and Metric was the observed variable. Comparable analyses were not conducted in younger adults due to their restricted age range. Results are presented in Table 5 and Figure 3.

**Figure 3.**
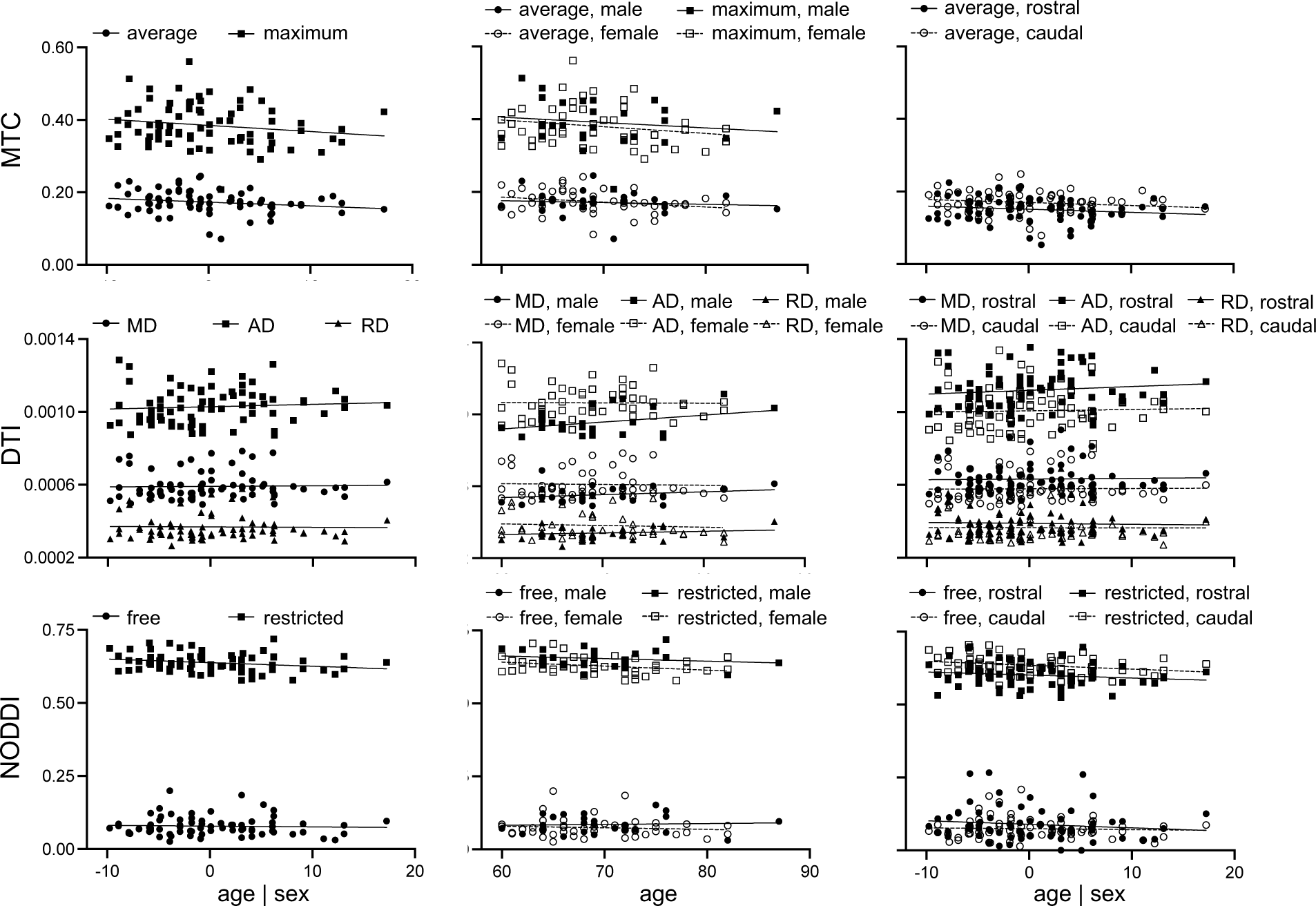
Relationships between chronological age and locus coeruleus structure within older adults are shown for each imaging metric (left column), with additional breakdowns by sex (middle column) and locus coeruleus subdivision (right column).

**Table 5.**
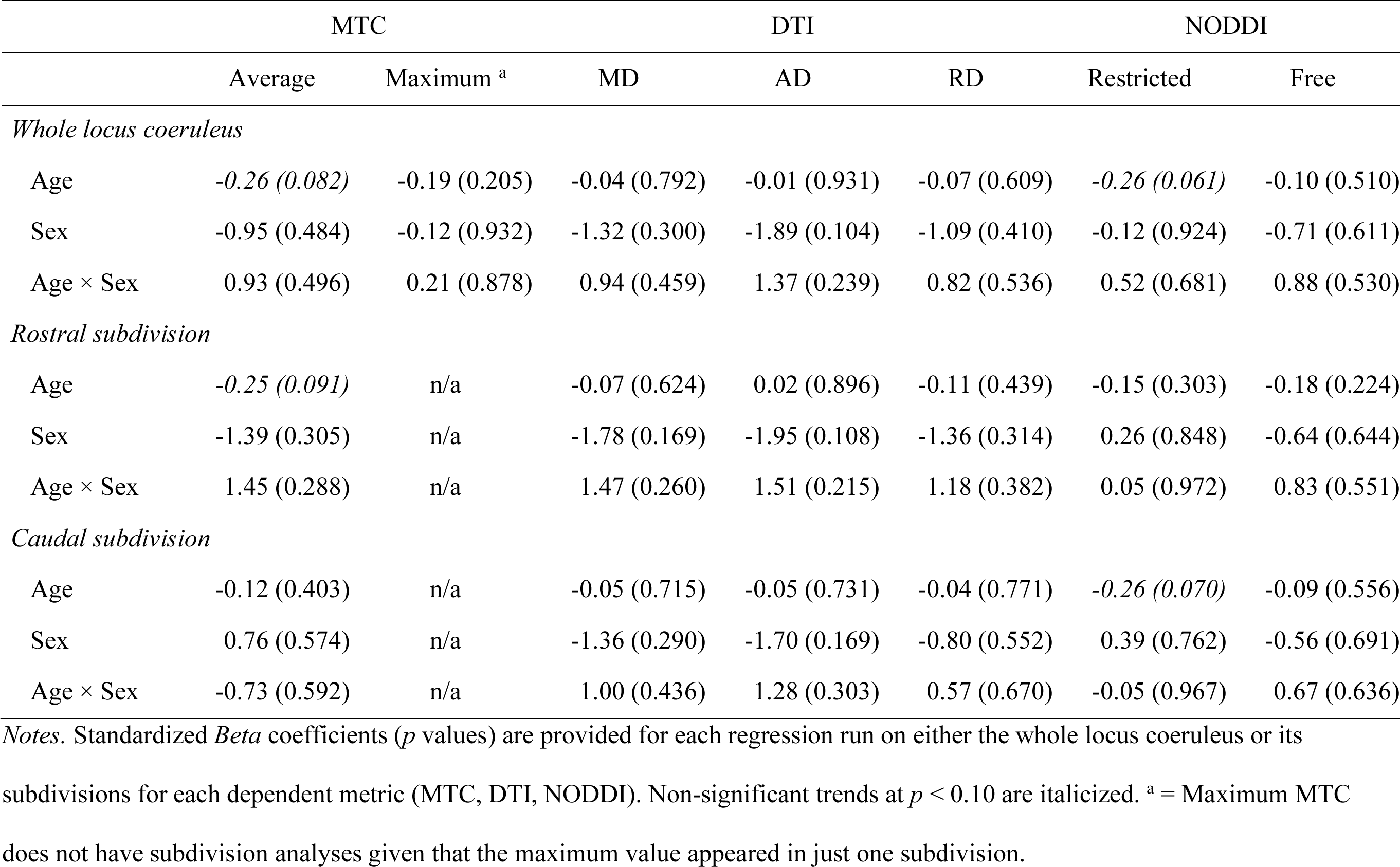
Relationships to Age Within Older Adults.

The whole locus coeruleus analyses revealed non-significant trends for older age relating to lower average MTC and lower restricted diffusion. The locus coeruleus subdivision analyses further revealed that the non-significant trends were seen in the rostral subdivision for average MTC and caudal subdivision for restricted diffusion. However, no effects of Age, Sex, or their interaction were statistically significant.

### Locus Coeruleus Structure Relates to Memory Performance

In all participants, separate partial correlations were conducted between each memory metric (mean recall, recall variability) and each metric from each imaging modality (MTC: average, maximum; DTI: MD, AD, RD; NODDI: restricted, free) in the whole locus coeruleus, controlling for age and sex. Significant effects survived Bonferroni correction for two (MTC, NODDI; *p* < 0.025) or three (DTI; *p* < 0.016) comparisons for each “family” of imaging metrics per memory metric. Results are presented in Table 6 and Figure 4.

**Figure 4.**
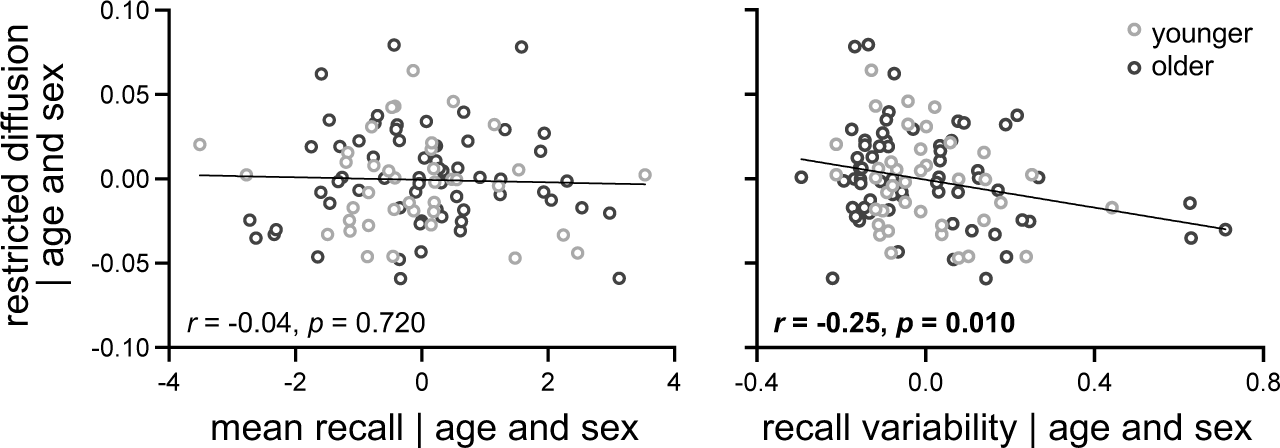
Relationships between memory performance (mean recall, recall variability) and restricted diffusion in whole locus coeruleus are shown, controlling for age and sex.

**Table 6.** Locus Coeruleus Structure Relates to Memory Performance.

|  | Mean Recall | Recall Variability |
| --- | --- | --- |
| *MTC metrics* |  |  |
| Average | 0.19 (0.054) | -0.13 (0.193) |
| Maximum | 0.15 (0.124) | -0.13 (0.165) |
| *DTI metrics* |  |  |
| MD | >-0.01 (0.983) | *0.20 (0.035)* |
| AD | 0.12 (0.232) | 0.12 (0.217) |
| RD | -0.04 (0.653) | 0.16 (0.100) |
| *NODDI metrics* |  |  |
| Restricted | -0.04 (0.720) | **-0.25 (0.010)** |
| Free | 0.07 (0.458) | -0.05 (0.587) |

*Notes.* Pearson $R$ statistics ( $p$ values) are provided for each partial correlation between memory performance (mean, variability) and each metric from the whole locus coeruleus. Significant Bonferroni corrected effects (MTC, NODDI: $p < 0.025$ ; DTI: $p < 0.016$ ) are bolded and non-significant trends ( $p < 0.05$ ) are italicized.

Results revealed that better (lower) recall variability was significantly related to higher restricted diffusion. Follow-up partial correlations in each locus coeruleus subdivision revealed that this relationship was significant for the caudal, *r* = -0.22, *p* = 0.025, but not rostral, *r* = -0.17, *p* = 0.087, subdivision. Follow-up partial correlations in each age group revealed that this relationship was significant for older, *r* = -0.26, *p* = 0.040, but not younger, *r* = -0.26, *p* = 0.115, adults.

### Relationships Among Locus Coeruleus Structure Metrics

In all participants, separate partial correlations were conducted among all metrics from each imaging modality (MTC: average, maximum; DTI: MD, AD, RD; NODDI: restricted, free) in the whole locus coeruleus, controlling for age and sex. Significant effects survived Bonferroni correction for six comparisons for each metric (*p* < 0.008). Results are presented in Table 7.

**Table 7.** Correlation Matrix for Locus Coeruleus Structure Metrics.

|  | Average | Maximum | MD | AD | RD | Restricted |
| --- | --- | --- | --- | --- | --- | --- |
| *MTC metrics* |  |  |  |  |  |  |
| Average |  |  |  |  |  |  |
| Maximum | **0.41 (<0.001)** |  |  |  |  |  |
| *DTI metrics* |  |  |  |  |  |  |
| MD | -0.13 (0.159) | -0.02 (0.863) |  |  |  |  |
| AD | -0.11 (0.221) | 0.04 (0.685) | **0.86 (<0.001)** |  |  |  |
| RD | -0.13 (0.174) | -0.12 (0.202) | **0.77 (<0.001)** | **0.58 (<0.001)** |  |  |
| *NODDI metrics* |  |  |  |  |  |  |
| Restricted | 0.13 (0.187) | 0.12 (0.202) | **-0.379 (0.001)** | **-0.388 (<0.001)** | *-0.234 (0.013)* |  |
| Free | -0.02 (0.813) | 0.12 (0.221) | 0.12 (0.200) | 0.02 (0.861) | 0.14 (0.137) | **0.25 (0.008)** |

*Notes.* Pearson *R* statistics (*p* values) are provided for each partial correlation between each metric from the whole locus coeruleus.
Significant Bonferroni corrected effects ( $p < 0.008$ ) are bolded and non-significant trends ( $p < 0.05$ ) are italicized.

Results revealed significant positive relationships between metrics within each imaging modality. Lower restricted diffusion was also significantly related to higher MD and AD. Follow-up partial correlations within each locus coeruleus subdivision revealed that the relationships were significant in both caudal and rostral subdivisions, except for relationships between restricted and free diffusion, *p*s > 0.11. Follow-up partial correlations within each age group revealed that the relationships were significant in younger and older adults.

## Discussion

The current study examined effects of age on locus coeruleus structure using a combination of magnetization transfer- and diffusion-weighted MRI in younger and older adults. Our approach extended prior work by examining age effects between younger and older adults as well as within older adults; including both MTC and DTI metrics and, for the first time, reporting NODDI metrics of locus coeruleus structure; and accounting for sex in all analyses. We were further interested in whether individual differences in locus coeruleus structure related to variability in memory performance. Key findings include that (1) maximum MTC in the locus coeruleus was more sensitive to differences between younger and older adults than average MTC, (2) DTI and NODDI metrics also showed significant, albeit smaller, age group differences, (3) males showed larger age group differences in MTC and NODDI metrics than females, (4) larger age group differences were seen in both rostral (DTI) and caudal (NODDI) subdivisions of the locus coeruleus, (5) however, no metrics of locus coeruleus structure were sensitive to effects of age within older adults, and (6) independent of age and sex, locus coeruleus structure (NODDI) related to variability in, but not mean, recall performance. These novel contributions to our understanding of the aging of locus coeruleus structure are detailed below.

When comparing extreme age groups, significantly higher MTC was seen in older than younger adults for both the maximum and average metrics, comparable to at least some prior studies (Clewett et al., 2016; Pievani et al., 2020). This finding is also consistent with there being a loss or reduction in locus coeruleus neurons with age (Flood & Coleman, 1988; German et al., 1988; Manaye et al., 1995; Vijayashankar & Brody, 1979) c.f. (Mouton et al., 1994). Other studies that found no age group differences in locus coeruleus signal intensity have used a combination of maximum (Dahl, Bachman, et al., 2023; Dahl et al., 2019) and average (Hämmerer et al., 2018; Porat et al., 2022) MTC, suggesting that metric type alone does not account for their discrepancies. We further showed, for the first time, that the age group difference was largest for the maximum relative to the average MTC metric, suggesting that future studies may improve their sensitivity to age group differences by using maximum MTC.

Relative to MTC, substantially fewer studies have examined aging of locus coeruleus structure using diffusion-weighted MRI, all of which modeled diffusion in each voxel as a single tensor. When using DTI, we observed significantly higher diffusivity in the locus coeruleus in older than younger adults and this age group difference was largest for AD relative to MD, but not significant for RD. Our age effects replicate at least one study that only looked at MD and similarly showed higher diffusivity in older than younger adults (Pievani et al., 2020), although it is inconsistent with prior work from our group that showed higher diffusivity in younger than older adults across all DTI diffusivity metrics (Langley et al., 2020, 2022) and another group that only looked at MD and RD (Porat et al., 2022). One explanation for this discrepancy is that our prior studies did not take sex into account, as was done here. Although Porat et al. considered sex and nonetheless found higher diffusivity in younger than older adults, albeit in some locus coeruleus hemispheres but not others across two datasets. Another, potentially more consequential, explanation is that the current study used a larger and less isotropic voxel (1.15×1.15×2.5 mm^3^) relative to prior work (Langley et al.: 0.95×0.95×1.0 mm^3^; Porat et al.: 1.7 mm^3^) to increase signal-to-noise for NODDI fitting. Whereas the non-isotropic voxel should not affect the diffusivity measures as they are directionally invariant, we may be more susceptible to partial volume effects with adjacent white matter (superior cerebellar peduncle) and the fourth ventricle.

Prior diffusion-weighted MRI studies that examined effects of age on locus coeruleus structure had not modeled diffusion using multi-compartment approaches that may better capture microstructural properties in gray matter as they estimate compartments of diffusion that are tissue invariant. When using NODDI, we observed significantly higher free diffusion in the locus coeruleus in older than younger adults, but the age group difference was not significant for restricted diffusion. Interestingly, whereas maximum MTC in the locus coeruleus was the single best predictor of age group, classification accuracy was slightly improved when adding DTI metrics to the model, but not when adding NODDI metrics. This suggests that NODDI may be less sensitive to aging of the locus coeruleus than DTI diffusivity, which contradicts what has been observed in other gray matter structures, such as the hippocampus (Venkatesh et al., 2020). Another possibility is that we may have too low signal-to-noise ratios for such complex modelling in this deep brain structure, as some participants had to be excluded from the NODDI analysis because of issues with model fitting (i.e., free diffusion values of zero). Nonetheless, finding that DTI and NODDI metrics show significant, albeit smaller, age group differences, and that DTI diffusivity improves the ability of maximum MTC to predict age group, indicates that diffusion metrics should be considered in future studies of locus coeruleus aging.

Of the previous aging studies that also examined sex difference in locus coeruleus structure, all were focused on MTC, and only one found lower maximum MTC in females than males that was independent of age group (Clewett et al., 2016). Most other studies either did not test for sex differences (e.g., (Dahl et al., 2019; Eckert et al., 2023) or reported no significant sex effects (Calarco et al., 2022; Jacobs et al., 2021; Liu et al., 2019; Shibata et al., 2006). In contrast, we observed significant effects of sex for all imaging modalities, with males having lower average MTC, lower DTI diffusivity, and higher NODDI diffusion relative to females. We further found interactions between age group and sex for average MTC and NODDI. That is, average MTC was significantly higher in older than younger males, but not females; whereas there were non-significant trends for lower restricted and higher free diffusion in older than younger males, but these effects did not approach significance in females. Given that these interactions between age and sex have been underreported, we note the importance for future studies to consider sex as a biological variable when examining effects of aging on locus coeruleus structure.

Whereas prior work using MTC in the locus coeruleus indicated that age effects may be more prominent in the rostral than caudal subdivision (Betts et al., 2017; Dahl et al., 2019; Liu et al., 2019, 2020), we did not find that the difference in average MTC between younger and older adults varied between the locus coeruleus subdivisions. This discrepancy may be due to differences in the way locus coeruleus is subdivided across studies, as it has been suggested that the rostral/caudal subdivisions may be overly simplistic and not reliably captured with current *in vivo* imaging approaches (Poe et al., 2020; Veréb et al., 2023). However, prior work finding significant effects have used both two (Dahl et al., 2019; Liu et al., 2019, 2020) and three (e.g., (Betts et al., 2017) subdivisions, indicating that the number of subdivisions alone cannot explain these findings. In line with this prior work, and with the notion that the rostral subdivision is more vulnerable to cell loss and reductions in cell density in aging, comparable to what is seen in Alzheimer’s Disease (German et al., 1992), we did find an interaction between age group and subdivision for the DTI metrics, with the age group difference in AD (higher in older than younger adults) being larger in the rostral than caudal subdivision. Although this finding contradicts our prior work showing comparable age group differences in locus coeruleus diffusivity in both subdivisions (Langley et al., 2022), we previously used a much smaller sample (35 younger, 28 older) and did not account for sex. The current study further found that the age group difference in NODDI diffusivity (higher in older than younger adults) was larger in the caudal than rostral subdivision, and that this interaction was driven by females relative to males. Thus, the different imaging modalities may be differentially sensitive to aging of locus structure across its subdivisions and interactions among these variables, as well as sex, warrants further investigation.

In contrast to the aforementioned results comparing younger and older adults, we did not find any significant relationships between chronological age and any measure of locus coeruleus structure within older adults. There were non-significant trends for older age relating to lower average MTC and lower restricted diffusion. The former is consistent with at least some prior studies (Calarco et al., 2022; Giorgi et al., 2022), and in the same direction as others that observed a decrease in locus coeruleus MTC with age after ∼60 years old (Eckert et al., 2023; Jacobs et al., 2021; Liu et al., 2019; Shibata et al., 2006). Looking across our extreme age group and within older adults findings for MTC, they are generally consistent with prior work reporting an inverted U-shaped function for locus coeruleus structure across the adult lifespan. However, age effects were larger between younger and older adults for maximum MTC but within older adults for average MTC, suggesting that future studies may improve their sensitivity to age effects by selecting the appropriate metric give their sample.

Independent of age and sex, we found that higher NODDI restricted diffusion in the locus coeruleus was significantly related to better (lower) recall variability, but not mean recall. Non-significant trends were also seen between higher average MTC and better mean recall, consistent with prior studies in older adults (Dahl et al., 2019; Dahl, Kulesza, et al., 2023; Elman et al., 2021; Hämmerer et al., 2018; Jacobs et al., 2021; Solders et al., 2020); and between higher MD and worse (higher) recall variability. Although the latter did not replicate our previous finding of higher DTI diffusivity relating to better mean recall within older adults (Langley et al., 2020, 2022), it does extend prior work by demonstrating the sensitivity of NODDI metrics in the locus coeruleus to memory performance, and the sensitivity of variability in memory performance to locus coeruleus structure. Our finding is also consistent with the notion that locus coeruleus structure contributes to its function via noradrenergic signaling. Individual and age-related differences in locus coeruleus structure may influence moment-by-moment attention states that, in turn, affect consistency in (variability) task performance, as previously shown for a working memory task (Unsworth & Robison, 2017). Recent animal studies have also demonstrated that a specific loss of locus coeruleus noradrenergic neurons was associated with worse memory performance (Gargano et al., 2023). Of note, our findings suggest that variability between trials may be more sensitive to degradation of locus coeruleus structure than average performance metrics (e.g., mean across trials) and should be considered in future work.

The current study aimed to bridge literatures that have examined age effects on locus coeruleus structure using different MRI modalities (magnetization transfer-weighted, diffusion-weighted) in different samples (between younger and older adults, within older adults) to identify which metrics are most sensitive to age and memory performance. When examining age effects between younger and older adults, we show that maximum MTC is the best predictor of age group, outperforming average MTC and both DTI and NODDI metrics that show significant, but smaller age effects. We further show that age group differences in locus coeruleus structure vary with sex and locus coeruleus subdivision, and we encourage future studies to consider their contributions. Within older adults, however, there were no significant effects of age on any measure of locus coeruleus structure. Finally, we demonstrated that individual differences in locus coeruleus structure (NODDI restricted diffusion) significantly relates to memory performance, and that variability in performance may be as, if not more sensitive than mean performance.

